# Haploid Asexual Blastocyst Fitness Varies Across Mouse Strains Related to Efficiency of Exit From Totipotency

**DOI:** 10.1101/2024.08.19.608531

**Authors:** Daphne Norma Crasta, Si Won Lee, Jolene Fredrickson, Theodore Thejo, Satish K. Adiga, Yulian Zhao, Guruprasad Kalthur, Nagarajan Kannan

**Affiliations:** Division of Reproductive Biology, Department of Reproductive Science, Kasturba Medical College, Manipal Academy of Higher Education, Manipal 576104, India; Division of Experimental Pathology and Laboratory Medicine, Department of Laboratory Medicine and Pathology, Mayo Clinic, Rochester, MN 55905, USA; Division of Reproductive Endocrinology and Infertility, Department of Obstetrics and Gynecology, Mayo Clinic, Rochester, MN 55905, USA; Centre of Excellence in Clinical Embryology, Department of Reproductive Science, Kasturba Medical College, Manipal Academy of Higher Education, Manipal 576104, India; Mayo Clinic Comprehensive Cancer Center, Mayo Clinic, Rochester, MN 55905, USA; Center for Regenerative Biotherapeutics, Mayo Clinic, Rochester, MN 55905, USA

**Keywords:** Parthenogenetic embryos, time-lapse imaging, first cleavage dynamics, totipotency clock, FVB mice, haploid blastocyst efficiency, asexual reproductive fitness, reproductive evolution, blastocyst, pre-implantation embryo development

## Abstract

*In vitro* activation, both sexually and asexually, facilitates assessing the reproductive mode and fitness of mammalian oocytes. Herein, we present evidence of the enhancement of asexual haploid blastocyst fitness in one selectively-inbred *Mus musculus* population. We tracked sexually and asexually activated-oocytes as they exited totipotency and self-organized into blastocyst-stage embryos. We examined haploid and diploid parthenogenetic potential of activated-oocytes. Unexpectedly, ∼90% of selectively-inbred mouse oocytes that were asexually activated successfully generated haploid blastocysts, contrasting with ∼90% failure in randomly-outbred mice. Furthermore, by closely tracking the timeline of exit from totipotency, we propose a novel ‘self-correcting’ ‘totipotency clock’, crucial for timely exit from totipotency and successful embryogenesis across mammals. Insufficiency in this ‘self-correcting’ prerequisite, will alter the fitness landscape in different reproductive modes. Collectively, this work provides a quantitative framework to investigate the unknown disruptive evolutionary trajectories of reproductive modes and fitness of females in anisogamous species.

**Highlights:**

- Serendipitious discovery of disruptive evolution of haploid asexual reproductive mode and preimplantation embryogenetic fitness in FVB strain of mice.
- Novel self-correcting totipotency clock regulates blastulation potential in mammals including humans and limits haploid asexual embryogenesis
- Evolution of haploid asexual reproductive mode and preimplantation embryogenetic fitness in FVB mouse is linked to a superior self-correcting totipotency clock lacking in other animals.

**Graphical Summary:** 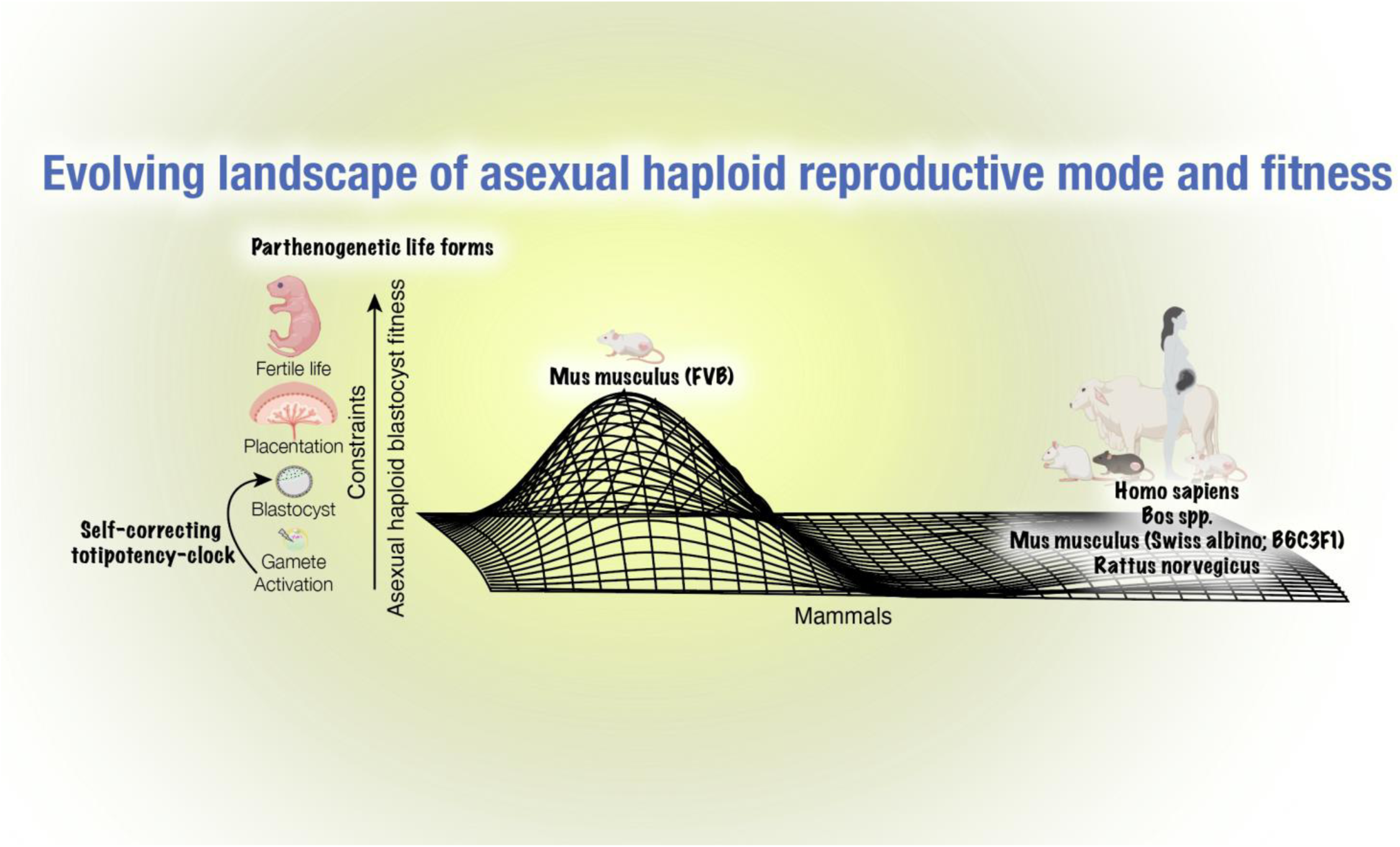

## Introduction

Charles Bonnet’s seminal discovery of parthenogenesis in 1740, demonstrating that spindle tree female aphids reproduced without male contribution, laid the groundwork for understanding the roles of sex in reproduction and evolution (Bonnet, 1745). Parthenogenesis, the clonal asexual generation by females, is widespread across invertebrates (Edwards et al., 2003; Lourenço, 2008; Suomalainen, 1950) and cold-blooded vertebrates (Booth et al., 2012; Watts et al., 2006; Wright and Vitt, 1993), occurring in species with exclusive asexual (Innes and Hebert, 1988; Suomalainen et al., 1987) or sexual (Braga-Pereira and Santos, 2021) reproduction and those capable of switching between the two modes (Cook, 1993; Grant, 1978; Liegeois et al., 2021; Puterka et al., 2012; Sperling et al., 2023; Suomalainen, 1950; Watts et al., 2006). Even within a species, distinct populations may adopt divergent reproductive strategies, highlighting the adaptability of reproductive modes (Snyman and Xu, 2023; Wright and Vitt, 1993). Notably, parthenogenesis has been observed in warm-blooded animals such as birds in captivity (Oellacher, 1872; Olsen, 1975; Parker and McDaniel, 2009) and sharks (Chapman et al., 2007; Dudgeon et al., 2017; Holtcamp, 2009).

In mammals, there are no documented instances of natural parthenogenesis yielding viable offspring, although development *in vivo* up to post-implantation embryo stages have been reported in mice (Kaufman and Howlett, 1986; Kaufman et al., 1977), suggesting that mammalian evolution may be a product of condition that selected binary sex roles while heavily suppressing asexual reproduction. Some species, such as certain lizard populations, have transitioned from sexual to exclusively asexual reproduction mode (Freitas et al., 2016), a phenomenon not seen in mammals. Several anecdotal reports exist on lone female birds (Oellacher, 1872; Olsen, 1975; Parker and McDaniel, 2009) and reptiles (Booth et al., 2023; Booth et al., 2012; Card et al., 2021; Watts et al., 2006) in captivity, to have exhibited unusual parthenogenetic capability, a phenomenon never observed in their counterparts living under natural habitats. The oocyte activation necessary for parthenogenesis involves a wide range of mechanisms, spanning from microbial endosymbionts to factors affecting calcium efflux. These activations occured unintentionally through diverse sources found in the environment or within an organism’s own physiology (Ma and Schwander, 2017).

Dr. Gregory Pincus and colleagues in 1936 conducted the first laboratory experiment demonstrating that parthenogenesis can be artificially induced in mammals (Pincus, 1936). Using artificial methods to activate rabbit oocytes, they successfully induced diploid parthenogenetic development (Pincus and Shapiro, 1940), paving the way for similar experiments in other mammalian species like mice, rats, pigs, horse, sheep, goats, etc (Kharche and Birade, 2013; Paffoni et al., 2008). Expansion of artificial methods (chemical, temperature, electrical) made it increasingly clear that parthenogenetic oocyte activation is not the major evolutionary barrier in emergence of parthenogenetic mammals in nature (Mitalipov et al., 2001; Paffoni et al., 2008). The activation using chemical agent like strontium (Sr^2+^) successfully mimicks the sperm-induced calcium oscillations, and is widely used in mice oocyte parthenogenetic activation (Liu et al., 2002). However, strontium is debated to be unable to cause the required calcium release in other mammalian oocytes like human, pig or cow (Swann, 2018), and alternate activating agents are used in these species. Nevertheless, when these parthenogenetic embryos were transferred into surrogate mothers, they did not grow to term, as observed in mice [day 10; (Surani and Barton, 1983)], rabbits [day 11; (Ozil, 1990)], sheep [day 21; (Loi et al., 1998)], pigs [day 30; (Kawarasaki et al., 2009)] and goats [day 34; (Kharche et al., 2014)]. Similarly, oocytes parthenogenetically activated *in vivo* also get blocked at various stages of post-implantation embryonic development (Hao et al., 2006; Taylor, 1996), indicating that male gamete factors are required to overcome this reproductive constraint. This is attributed to genomic imprinting, an epigenetic process that silences one allele based on its parental origin, leading to monoallelic expression (Barton et al., 1984; McGrath and Solter, 1984; Surani et al., 1984). Imprint marks are inherited by daughter cells but are erased and re-established during gametogenesis (Reik et al., 2001). Paternally expressed genes like *Igf2*, *Mest*, and *Peg3* promote fetal growth, while maternally expressed genes like *Igf2r*, *H19*, and *Grb10* restrict it (Charalambous et al., 2003; Lau et al., 1994; Lefebvre et al., 1998; Leighton et al., 1995; Li et al., 1999). In this direction, the implantation of bi-maternal parthenogenetic embryos obtained from reconstructed oocytes with experimental manipulation of the *Igf2/H19* and *Dlk1/Gtl2* (MEG3) imprinted loci using mutant mice, successfully enabled the production of live and reproductively fit mice (Kono et al., 2004), which highlight that the developmental constraints of parthenogenetic embryos are primarily due to epigenetic factors (Kono et al., 2002). Human oocytes can undergo parthenogenetic activation *in vitro* (de Fried et al., 2008; Gook et al., 1995; Osman et al., 2019; Van Blerkom et al., 1994), but their further development is restricted at the 2-4 cell stage (Taylor, 1996).

Selective inbreeding for ‘desirable’ traits in animals and plants artificially shapes the genetic makeup of populations, rapidly altering timelines compared to natural evolution (Brotherstone and Goddard, 2005; Hill, 1982; Hill and Kirkpatrick, 2010). However, the impact of trait-selective inbreeding pressure on reproductive evolutionary trajectories remains poorly understood in mammals.

Asexual or sexual mode of oocyte activation under *in vitro* conditions provides crucial insights into the fitness landscape of preimplantation embryonic development (Crasta et al., 2023; Liu et al., 2002). The mammalian activated oocyte differentiates into a blastocyst within 5-7 divisions (Hardy et al., 1989; Niwayama et al., 2019). In this study, we examined sexually (sperm-activated) and asexually (parthenogenetic) activated oocytes from both outbred and inbred mouse (*Mus musculus*) strains. Utilizing time-lapse imaging, we tracked the initial totipotent cell, i.e., the activated oocyte, and its symmetrical self-renewal division, giving rise to two totipotent blastomeres. Subsequently, we tracked their exit from the totipotent-state, observed as the transition from 2-cell-to-4-cell embryo (Guo et al., 2019), along with an additional 3-5 divisions, during which blastomeres differentiated into pluripotent embryonic and extraembryonic components of a progressively developing blastocyst. Our results reveal a previously unobserved remarkable enhancement in haploid parthenogenetic fitness of activated oocytes in an inbred (FVB) strain of mice, marking the first evidence of controlled disruptive evolution in mammalian reproductive modes and fitness landscape. Additionally, we identified a novel ‘self-correcting’ earliest developmental clock operating to ensure activated oocytes, irrespective of reproductive mode or species, exit totipotency ‘on-time’. This provides a robust universal model for assessing sexual and asexual mammalian blastocyst generation efficiency and quality, with implications extending to human reproduction.

## Results

### Tracking Oocyte Reproductive Modes Unearths Divergent Oocyte Asexual Embryogenetic Fitness Landscape in *Mus musculus* Populations

Mice used in the present study are descendants of an initial outbred *Mus musculus* population, which can be traced back to the Swiss mice established in 1932 at Rockefeller Institute in USA (Rice and O’Brien, 1980). While the general research purpose Swiss albino population arose in the laboratory from randomly outbred Swiss mice, certain Swiss mice underwent two consecutive rounds of selective-breeding. Initially, they were inbred for traits associated with sensitivity to histamine diphosphate, followed by selection for the presence of the <u>F</u>riend leukemia <u>v</u>irus, <u>B</u> strain resulting in the development of the FVB population (Cui et al., 1993; Rice and O’Brien, 1980).

To understand reproductive modes and blastocyst fitness of *Mus musculus*, we investigated three model systems: *in vitro* fertilization (IVF)-derived <u>sex</u>ually fertilized <u>e</u>mbryos (**SexE**) and <u>asex</u>ually fertilized <u>e</u>mbryos (ASexE), generated using strontium chloride activation in the presence and absence of cytochalasin D to obtain <u>dip</u>loid (**Dip-AsexE**) and <u>hap</u>loid asexual parthenogenetic embryos (**Hap-AsexE**), respectively (schematics in **Figure 1A**). Ploidy status of Hap-AsexE and Dip-AsexE was confirmed through metaphase-karyotyping (data not shown). The comparable efficiency of oocyte activation in both *Mus musculus* populations suggest a remarkable similarity in the developmental competence of their oocytes (**Supplementary Figure 1A**). A comprehensive *in vitro* imaging analysis was conducted on 608 activated oocytes, examining their developmental trajectories as they exited totipotency (Vega-Sendino et al., 2024) and self-organized into blastocyst-stage embryos under both sexual and asexual reproductive conditions (**Figure 1B-C**; **Supplementary Figure 1B-C**).

**Figure 1:**
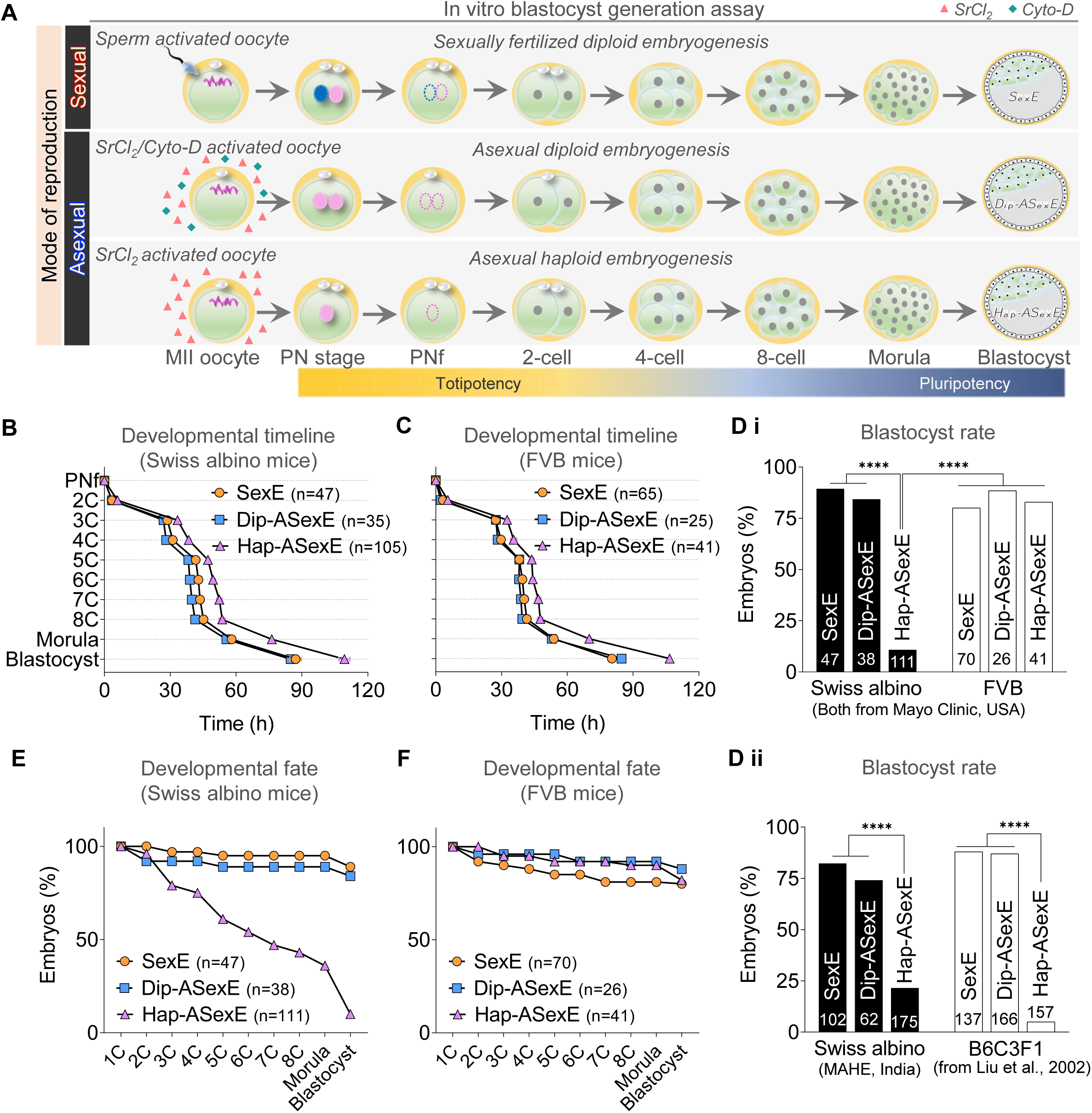
Assessment of Sexual and Asexual Reproductive Mode and Blastocyst Fitness in inbred and outbred *Mus musculus* populations. **A)** Schematic representation of blastocyst generation assay through *in vitro* fertilization and parthenogenetic activation using SrCl_2_ with and without cytochalasin D, showcasing ability of oocytes to progress to the blastocyst-stage. **B)** Developmental timelines from pronuclei fading (PNf) to blastocyst-stages of sexually fertilized (SexE), diploid parthenogenetic (Dip-AsexE), and haploid parthenogenetic (Hap-ASexE) embryos from Swiss albino strain of mice. **C)** Developmental timelines from PNf to blastocyst-stages of SexE, Dip-AsexE, and Hap-ASexE from FVB strain of mice. **D)** Blastocyst rates in SexE, Dip-AsexE, and Hap-ASexE of **i)** Swiss albino and FVB mice from Mayo Clinic, USA, **ii)** Swiss albino from MAHE, India and B6C3F1 (Liu et al., 2002). The groups were compared using Fisher’s exact test, **** p<0.0001. **E)** Developmental fate of SexE, Dip-AsexE, and Hap-ASexE derived from Swiss albino strain of mice. **F)** Developmental fate of SexE, Dip-AsexE, and Hap-ASexE derived from FVB strain of mice. **See also, supplementary figure 1**.

From pronuclei fading (syngamy) to blastocyst, the preimplantation stages of embryo development unveiled marked delay in haploid parthenogenetic embryo (Hap-AsexE) development compared to the diploid embryos (SexE and Dip-ASexE) in both strains of mice (**Figure 1B-C**). Hap-AsexE displayed prolonged durations of the embryonic cell cycle (**Supplementary Figure 1D-G**), accompanied by delayed blastomere synchrony (**Supplementary Figure 1H-I**). The absence of this delay in Dip-AsexE strongly suggests that the presence of a single set of uniparental genetic components is responsible for the prolonged cell cycle durations and slower development (Henery and Kaufman, 1992).

All three groups of zygotes from both *Mus musculus* populations demonstrated equally high blastocyst development efficiency, with the exception of Hap-ASexE from the Swiss albino strain (**Figure 1Di**). The poor blastocyst rate in Swiss albino mice-derived Hap-ASexE was confirmed in two separate set of experiments in two different laboratories, located in different continents (**Figure 1Di and 1Dii**), and a similar observation has been reported by Liu et al., 2002 in B6C3F1 mice (**Figure 1Dii**). Detailed examination of the morphokinetics of Swiss albino-derived Hap-ASexE, showed regression in their development (**Figure 1E**), resulting in few embryos reaching the blastocyst stage with poor morphology and limited growth potential (**Supplementary Figure 1J**). In contrast, Hap-ASexE derived from FVB mice exhibited a developmental trajectory comparable to SexE and Dip-AsexE (**Figure 1F**), resulting in well-expanded blastocysts (Supplementary Figure 1J). Additionally, there were significant differences observed between Hap-ASexE from both the *Mus musculus* populations in terms of embryonic cell cycle durations and blastomere synchrony during these cycles (**Supplementary Figure 1F-I**).

Hence we report that, the diploid parthenogenetic efficiency approaches near perfection when compared to SexE, indicating minimal constraints in acquiring diploid asexual development. Conversely, Hap-AsexE appears to face significant constraints, except for FVB, which seems to have disruptively evolved to achieve comparable, if not slightly superior, blastocyst fitness compared to sexual reproduction. The rapid emergence of enhanced blastocyst fitness of oocyte’s haploid asexual reproductive mode in FVB mice inbred for specific phenotypes unrelated to reproduction (**Figure 2A**), suggests that reproductive modes and their corresponding blastocyst fitness levels might be diversifying more rapidly and in unpredictable ways.

**Figure 2:**
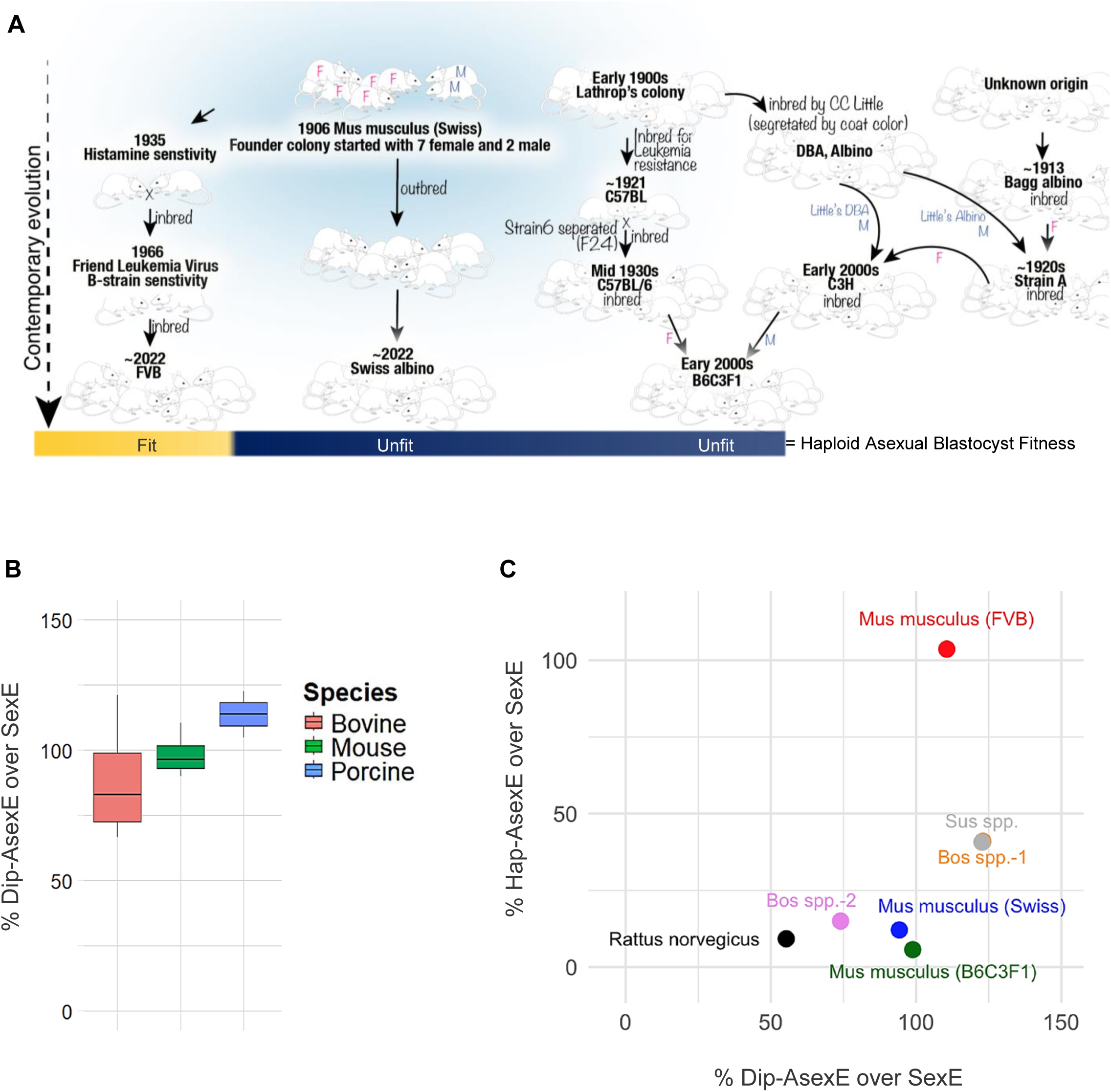
Cross-species and cross-strain examination of mammalian reproductive embryogenetic fitness. **A)** Contemporary Evolution of sub-strains of the *Mus musculus* (Swiss albino, FVB and B6C3F1 mice). Data obtained from (Cameron et al., 1985; Cui et al., 1993; Kern et al., 2012; Mekada et al., 2009; Rader, 2018; Rice and O’Brien, 1980), (https://www.cancerresearch.org/blog/november-2014/birth-of-the-lab-mouse) and (https://www.smithsonianmag.com/science-nature/history-breeding-mice-scienceleads-back-woman-barn-180968441/). **B)** Whisker plot depicting the variation in the developmental efficiencies of Dip-ASexE in comparison to SexE in different species. Data obtained from our study, as well as from (Cevik et al., 2009; Dinnyes et al., 2000; Gupta et al., 2009; Lagutina et al., 2004; Liu et al., 2002; Park et al., 2011; Wang et al., 2008). **See also, supplementary table 2**. **C)** Cross-species assessment of developmental efficiencies of Dip-ASexE and Hap-AsexE in comparison to SexE, showing enhanced blastocyst fitness of Hap-ASexE derived-from FVB strain of mice, compared to other mice stains (Swiss albino, B6C3F1) and species (*Rattus norvegicus, Bos spp., Sus spp.*). Data obtained from our study, as well as from (Lagutina et al., 2004; Liu et al., 2002; Park et al., 2011; Roh et al., 2003; Wang et al., 2008). **See also, supplementary table 1**.

### Cross-Species and Cross-Strain Examination Shows Uniquely Evolved Reproductive Advantage in Inbred FVB Strain of *Mus musculus*

Next, to understand cross-species and cross-strain differences, we surveyed literature and found 5 studies that reported Hap-ASexE, Dip-ASexE and SexE blastocyst rates in mammals including mice (Liu et al., 2002), rat (Roh et al., 2003), bovine (Lagutina et al., 2004; Wang et al., 2008) and porcine (Park et al., 2011) (**Supplementary Table 1**). The findings were striking. In all mammals, the Dip-AsexE efficiency approached near perfection when compared to SexE, despite the slight variation in blastocyst efficiencies of various studies for the same species (**Figure 2B, Supplementary Table 2**). Conversely, in all animals except FVB mice (present study), Hap-AsexE had highly constrained and poor developmental fate (**Figure 2C, Supplementary Table 1**).

FVB mice, originally derived through selective breeding for histamine sensitivity following pertussis vaccination, were inbred for eight generations to achieve homozygosity, resulting in the fixation of the *Fv-1^b^* allele associated with susceptibility to the B-strain of Friend leukemia virus (Kirczenow MacDonald et al., 2021)(**Figure 2A**). Whole-genome sequencing by Thomas Keane and colleagues at the Welcome Trust Sanger Center identified 286 FVB-specific non-synonymous SNPs, including previously known significant mutations in FVB in the *Pde6b* gene, linked to retinal degeneration, a 2-bp deletion in the *Hc* gene, impairing complement 5 secretion, and modifications in T-cell receptor variable regions related to resistance to collagen-induced arthritis (Wong et al., 2012). Additional private mutations were found in genes such as *Scn10a*, *Rnf186*, *Olfr1469*, *Foxa3*, *Krt5*, *Tpbg*, *Apbb1*, *Arhgef5*, *Ube2l6*, *Cpt1a*, *Rpl37-ps1*, *Ermap*, *Hmcn1*, *Olfr1454*, *Dtx4*, *Zfp521*, *Olfr922*, and *Dusp27* (Wong et al., 2012). Despite these insights, the genetic and molecular mechanisms underlying histamine sensitivity remain elusive, likely due to the polygenic and physiological intricacies of the trait. This genetic complexity, along with the presence of the *Fv-1^b^* allele and numerous other FVB-specific mutations, may contribute to the observed variation in the survival of haploid blastocysts in FVB mice, warranting further investigation to pinpoint the precise genetic factors involved.

### First Cleavage Dynamics of Self-renewing Totipotent Oocyte and its Embryogenetic Fitness in Mus musculus Populations

To further understand the divergent oocyte embryogenetic fitness observed above, we examined the oocytes as they divided into 2-cell totipotent blastomeres (Condic, 2014). The time-lapse imaging identified that the activated oocyte-to-2-cell (**2C**) embryo development progressed through dynamic first-cleavage (**FC**)-states (Ezoe et al., 2019). We refined the characterization of FC-states. Based on the observed irregularity during dynamic FC-states (from initial 2C appearance to proper 2C formation), we devised a scoring system ranging from FC-I to -IV. These FC-states were discerned using visible microscopic phenotypes, such as the initial appearance of the cleavage axis, subsequent irregular fragmentation, and eventual reappearance of the proper 2C cleavage axis. Therefore, the phases of totipotent zygote to 2-cell division encompass FC-states denoted as ‘early-2C’, ‘irregular-2C’, and ‘proper-2C’ phases of the first cleavage (**Figure 3A**).

**Figure 3:**
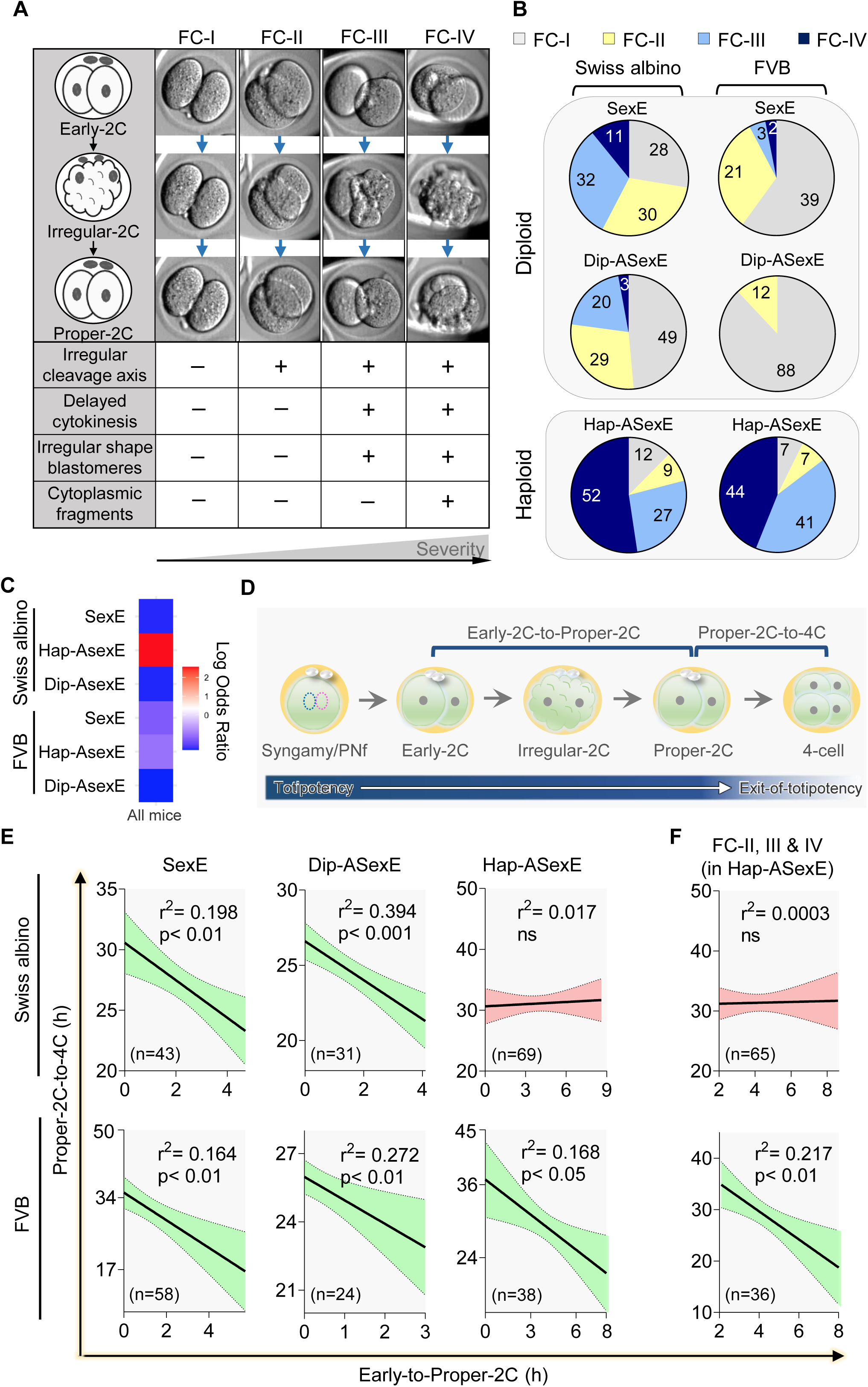
Symmetric Self-renewing Totipotent Cell First Cleavage Dynamics and Essential Role of Self-correcting Totipotency Clock in Sexual and Asexual Embryogenetic Fitness of *Mus musculus*. **A)** Illustration, representative images and scoring criteria for FC-I/II/III/IV during first-cleavage in mice embryos captured through time-lapse monitoring. **See also, supplementary videos 1-4**. **B)** Incidence of FC-states in SexE, Dip-AsexE and Hap-ASexE derived from Swiss albino and FVB strains of mice. **C)** Heat map showing Odds Ratio (OR) for probability of SexE, Dip-AsexE and Hap-ASexE derived from Swiss albino and FVB strains of mice to achieve developmental success to the blastocyst-stage (OR below 1) or to undergo arrest in earlier stages of preimplantation embryo development (OR above 1). The values are normalized to the mean success rate of all the mice embryos combined. **D)** Schematics of early embryo cleavage dynamics from sexual/asexual fertilization to the 4-cell-stage. The stages include syngamy/PN fading, early-2C-stage, irregular-2C-stage, proper-2C-stage and 4-cell-stage. **E)** Correlation of developmental time between FC-state (early-to-proper-2C) and exit of totipotency (proper-2C-to-4C) in SexE, Dip-AsexE and Hap-ASexE, derived from Swiss albino and FVB strains. **F)** Correlation of developmental time between FC-state (early-to-proper-2C) and exit of totipotency (proper-2C-to-4C) in FC-II, III and IV embryos derived from Swiss albino Hap-ASexE and FVB Hap-ASexE. Linear regression analysis was performed for data in panels E and F. p-value<0.05 was considered significant, and the graph was shaded green. p-values>0.05 were considered non-significant (ns) and the graphs was shaded red. **See also, supplementary figures 2 and 3**.

FC-I displayed no irregularity (**Supplementary Video 1**), while FC-II exhibited minimal blastomere movement at the cleavage-axis (**Supplementary Video 2**), commonly observed in pre-implantation embryo cleavage. FC-III presented delayed cytokinesis and irregularly shaped blastomeres, along with an irregular cleavage-axis (**Supplementary Video 3**). FC-IV involved the presence of cytoplasmic fragments in addition to the characteristics observed in FC-III (**Supplementary Video 4**).

This dynamic-state was exclusive to the first-cleavage during embryogenesis, and all four FC-states were observed across *Mus musculus* populations (**Supplementary Figure 2A**). Notably, Hap-ASexE from both Swiss albino and FVB strains, with divergent blastocyst fitness (**Figure 1Di**) showed similar FC-state distribution with significantly high combined occurence of FC-III and -IV (80-85%; **Figure 3B**), suggesting ploidy-status influences FC-dynamics during first embryonic self-renewal division. On the other hand, *Mus musuculus* population-specific differences in FC-states were observed in SexE and Dip-AsexE, with FC-III and -IV completely absent in FVB Dip-AsexE.

To understand the biological significance of these dynamic FC-states in altering the developmental fate of activated oocytes, we calculated FC-state associated risk for the embryos to developmentally get arrested before reaching the blastocyst-stage. As predicted, the risk for developmental arrest was highest for Hap-ASexE derived from Swiss albino strain (**Figure 3C**). Upon further stratification of embryos by FC-state, it became evident that Hap-ASexE from Swiss albino mice, irrespective of FC-state, displayed a consistently higher odds ratio (OR) for developmental arrest before reaching the blastocyst stage (**Supplementary Figure 2B**). SexE from FVB mice showed a higher risk of arrest with FC-III and -IV. The remaining embryo groups displayed relatively successful developmental probabilities.

These findings highlight that normal embryo development may be influenced by the dynamic states of the first cleavage in a context-specific manner.

### Novel Self-Correcting Totipotency Clock for Robust Blastocyst Generation in *Mus musculus* **populations**

Investigation into whether FC-states influence early embryo morphokinetics prompted a thorough analysis of time-lapse data (**Figure 3D**). The developmental timeline for resolution of FC-state irregularity (early-to-proper-2C) was in the order of FC IV>III>II>I, in SexE, as well as Dip-AsexE and Hap-ASexE (**Supplementary Figure 2C-E**) across *Mus musculus* populations. In contrast, the developmental timeline for progression from the proper-2C-to-4C-stage, exhibited almost opposite trends i.e., FC I>II>III>IV, in SexE, as well as Dip-AsexE and Hap-ASexE (**Supplementary Figure 2F-H**) across *Mus musculus* populations. Unexpectedly, durations of FC-state irregularity (early-to-proper-2C) and proper-2C-to-4C development were significantly inversely correlated for Hap-AsexE in FVB mice (r^2^=0.168, p<0.05) but not in Swiss albino mice (r^2^=0.017, p=ns) (**Figure 3E**). More strikingly, there was a significant inverse correlation observed in SexE (Swiss albino, r^2^=0.198, p<0.01; FVB, r^2^=0.164, p<0.01) and Dip-AsexE (Swiss albino, r^2^=0.394, p<0.001; FVB, r^2^=0.272, p<0.01) in both strains of mice (**Figure 3E**). This clearly suggested a developmental clock operating in the first two divisions and may be essential for normal progression towards blastocyst development.

To elucidate the crucial role of the developmental clock associated with early cleavage dynamics in driving normal embryo development *in vitro*, we investigated whether similar correlations exist for embryos classified as FC-II/III/IV states as FC-I showed no irregularities (**Figure 3F**). Our analysis revealed a significant inverse correlation in the hap-ASexE from FVB mice (r^2^=0.217, p<0.01) but not in Swiss albino mice (r^2^=0.0003, p=ns) (**Figure 3F**). This suggests that contemporary evolution has fundamentally altered the oocytes, enabling SexE and Dip-AsexE to execute a temporal balancing act to counter irregularities observed in the dynamic FC-states, in general, is extended to Hap-ASexE in FVB mice, whereas this phenomenon is not observed in Hap-ASexE in Swiss albino mice (**Figure 3F**). The SexE and Dip-AsexE across *Mus musculus* populations, and the Hap-AsexE embryos from FVB population that successfully formed a blastocyst, relied on the intricately maintained cleavage developmental clock (**Supplementary Figure 3Avi**), but this was not extended to the Swiss albino Hap-AsexE (**Supplementary Figure 3Aiii**). However, the embryos arrested in all groups across *Mus musculus* populations were due to a lack of maintenance of the cleavage developmental clock (**Supplementary Figure 3Ai-vi).**

The retrospective analysis of time for pronuclear fading (PNf) revealed that it was directly proportional to the FC-states (**Supplementary Figure 3B-D**) (Ezoe et al., 2019), while there was no association seen between the duration of PNf-to-early-2C-stage and the FC-states in any mice embryos (**Supplementary Figure 3E-G**).

### Totipotency Clock is also Essential for Oocyte Embryogenetic Fitness in *Homo sapiens* populations

To assess the applicability of the phenomenon observed in mice to other mammals, we conducted a retrospective analysis of time-lapse imaging data from human sexually fertilized embryos (Human-SexE) (n=93 human embryos) and haploid parthenogenetic embryos (Human-Hap-ASexE) (n=5 human embryos) obtained from the Mayo Clinic IVF Clinic. Utilizing the criteria outlined in **Figure 3A**, we characterized Human-SexE embryos and observed the presence of all four FC-states (**Figure 4A**; **Supplementary Videos 5 to 8**). Further, a small fraction of the total analyzed embryos (3.47%) were haploid (1PN) in nature (Human-Hap-ASexE), which showed only FC-states-II,-III and -IV (**Figure 4B**). The majority of Human-SexE (∼70%) exhibited FC-I and -II, with a lesser proportion (∼30%) displaying FC-III and -IV), while majority of Human-Hap-AsexE (80%) exhibited FC-III and -IV (**Figure 4B**), similar to the Hap-AsexE of Mus musculus populations (**Figure 3B**). The duration of early-to-proper-2C-stage increased with increase in FC-states (FC-IV>III>II>I; **Supplementary Figure 4A**), and showed opposite trend in duration of proper-2C-to-4C (**Supplementary Figure 4B**). Retrospective analysis of pronuclear fading (PNf) duration showed increase with increase in FC-states (**Supplementary Figure 4C**), while duration from PNf-to-early-2C-stage did not show any correlation (**Supplementary Figure 4D**). Most of the Human-Hap-ASexE arrested prior to first cleavage (99.56%), while the rest arrested after a few cell divisions (∼8-cell stage, 0.34%), with only 0.08% reaching the blastocyst stage. Assessment of human blastocyst quality in Human-SexE (Pierson et al., 2023), in relation to the FC-states at 2C-stage revealed intriguing findings. FC-I and -II embryos were associated with properly-differentiated inner cell mass and trophectoderm (**Figure 4C; Supplementary Figure 4E-F**), as indicated by low Gardner Grade-scores. In contrast, FC-III and -IV embryos were linked to poorly-differentiated inner cell mass and trophectoderm, exhibiting higher Gardner Grade-scores (Score 3 and above; **Figure 4C; Supplementary Figure 4E-F**). Furthermore, when correlating FC-states with clinical decisions in assisted reproduction, it was observed that Human-SexE with lower FC-states were more favorable for transfer to patients, whereas those with higher FC-states resulted in poor/ average-quality blastocysts deemed unsuitable for transfer (**Supplementary Figure 4G**).

**Figure 4:**
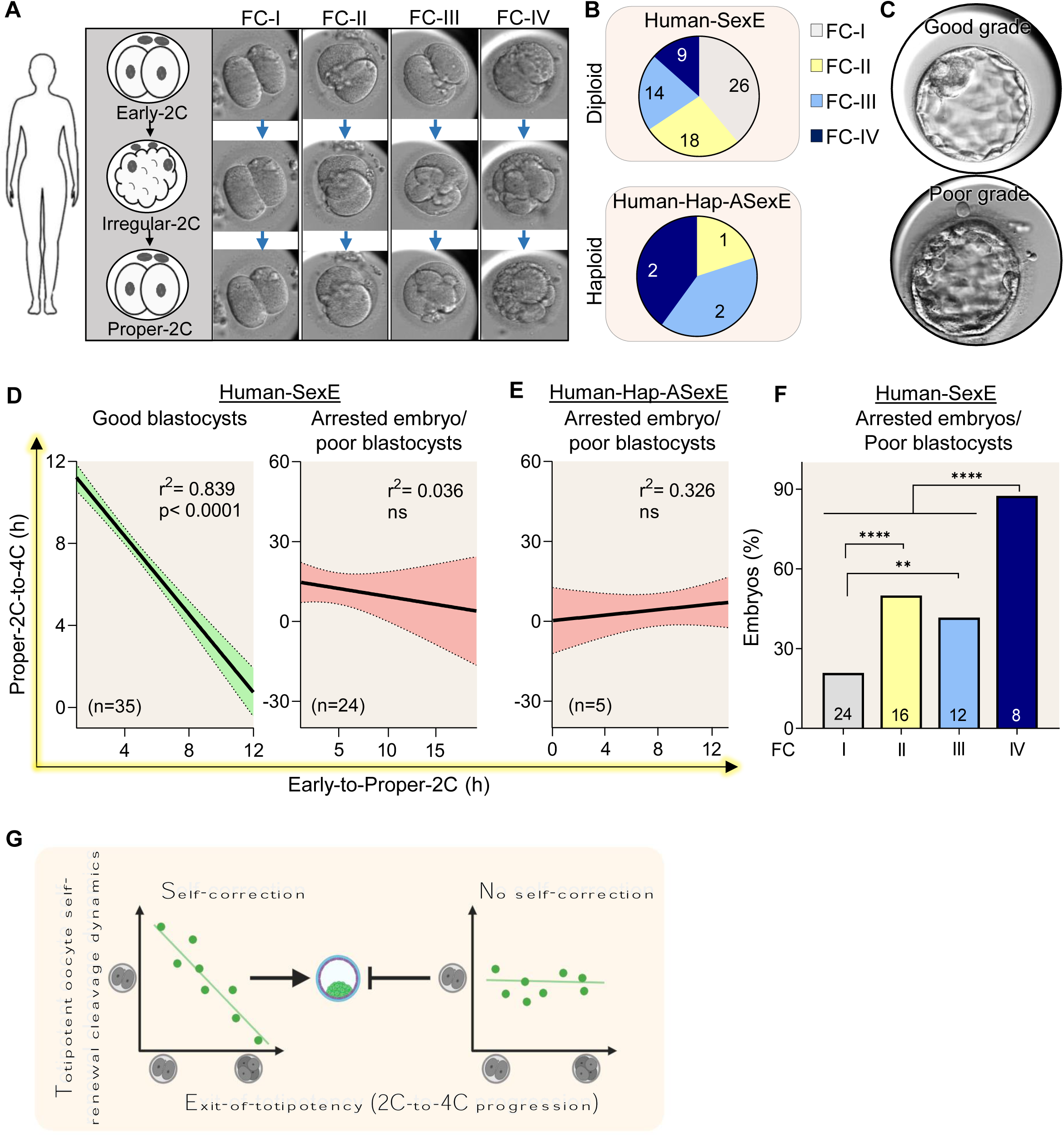
Essential Role of Self-correcting Totipotency Clock in Sexual Embryogenetic Fitness of *Homo sapiens*. **A)** Illustration and representative images for FC-states -I to -IV during first-cleavage in Human-SexE captured through time-lapse monitoring. **See also, supplementary videos 5-8**; **B)** Incidence of FC-states -I to -IV in Human-SexE and Human-Hap-ASexE. **C)** Representative images of good and poor Gardner grade human blastocysts. **D)** Correlation of developmental time between FC-state (early-to-proper-2C) and exit of totipotency (proper-2C-to-4C) in Human-SexE that formed good quality blastocysts, and embryos that got arrested or formed poor quality blastocysts. **E)** Correlation of developmental time between FC-state (early-to-proper-2C) and exit of totipotency (proper-2C-to-4C) in Human-Hap-ASexE that got arrested or formed poor quality blastocysts. Linear regression analysis of timelapse data performed for data in panels D and E. p-value<0.05 was considered significant, and the graph was shaded green. p-values>0.05 were considered non-significant (ns) and the graphs was shaded red. **F)** Effect of FC-states on the percentage of Human-SexE that got arrested or formed poor quality blastocysts. The data was analyzed using Fisher’s exact test, **** p<0.0001, ** p<0.01. **G)** Summary illustration showing the rescue mechanism in successful blastocyst forming embryos through subsequent cell cycle compensation, while failure to compensate leads to arrest or poor quality blastocysts. **See also, supplementary figure 4.**

Correlation analysis between the duration of FC-states and the duration of exit-from-totipotency in Human-SexE confirmed that embryos progressing to form good-quality blastocysts (fully expanded with properly-differentiated inner cell mass and trophectoderm) displayed a very strong negative correlation (r^2^=0.839, p<0.0001) (**Figure 4D; Supplementary Figure 4H-I**), indicative of temporal balancing act in the 2^nd^ embryonic cell cycle step, also observed in *Mus musculus* embryos (**Supplementary Figure 3A)**. This compensation, however, was not observed in arrested embryos or embryos resulting in poor-quality blastocysts derived from Human-SexE (r^2^=0.036, p=ns) (**Figure 4D; Supplementary Figure 4H-I**) or Human-Hap-ASexE (r^2^=0.326, p=ns) (**Figure 4E**), consistent with our observation in *Mus musculus* embryos (**Supplementary Figure 3A)**. No correlation was seen in the duration from early-to-proper-2C-stage versus the time of PNf (**Supplementary Figure 4J**). As expected, embryos exhibiting FC-IV were at the highest risk of developmental arrest or forming poor quality blastocysts (**Figure 4F**; Supplementary Figure 4K). Thus, the FC-states scoring system observed in the first-cleavage-stage of embryos, along with compensation in the subsequent cell cycle, can serve as an early predictive marker for blastocyst quality (**Figure 4G**), which is a crucial factor for success in assisted reproduction.

## Discussion

The cyclical nature of life’s reproduction hinges on the vital process of regeneration, underpinned by two fundamental mechanisms designed to perpetuate the existence of living systems. At its most primitive, this involves genome doubling and subsequent fission, yielding two compartments that faithfully recreate parental equivalent genomes in the offspring, a process often referred to as asexual or natural cloning. With further evolutionary refinement, this mechanism gave way to genome fission and fusion, a hallmark of sexual reproduction (Baker and Parker, 1973; Goodenough and Heitman, 2014). Within the context of multicellular organisms, particularly in mammals, the delineation between “male” and “female” assumes considerable biological significance in the landscape of sexual reproduction. These designations delineate specialized roles in gamete production, reproductive anatomy, and contribute significantly to evolutionary processes such as sexual selection and genetic diversity maintenance within populations (Cade, 1984; Dimijian, 2005; Dixson, 2021). Nature provides ample evidences that animals with sexual reproduction have evolved to adopt asexual modes of reproduction improving their embryogenetic fitness to conserve their species (Avise, 2008; Beukeboom and Vrijenhoek, 1998; Janko et al., 2023). We lack contemporary evidence of natural selective pressure widening reproductive fitness landscape of animals.

We present compelling evidence of significant enhancement in the reproductive modes and blastocyst fitness of oocyte totipotency, particularly in haploid parthenogenetic blastocyst developmental potential, within the FVB mouse population. This stands in stark contrast to the observed poor efficiency of haploid parthenogenesis in the outbred Swiss albino mice and the B6C3F1 mice (Liu et al., 2002). Studies have shown that these strains share genetic similarities with wild mouse populations, indicating their common ancestry with wild mice. However, extensive selective breeding in laboratory settings has led to specific genetic variations and adaptations unique to each strain. Genome sequencing and analysis have revealed the genetic diversity within and between mouse strains (Keane et al., 2011), shedding light on the genetic basis of various traits and susceptibility to diseases. This information is crucial for understanding the genetic mechanisms underlying complex traits and diseases, as well as mapping associated blastocyst fitness landscape. The genealogy of present-day inbred FVB with unique phenotypes including large pronuclear morphology (Wong et al., 2012) and outbred Swiss albino mouse (Mus musculus) strains can be traced back to seven female and two male mice imported from Switzerland to the United States in 1926 by Clara J. Lynch (Lynch, 1969). These mice were subsequently distributed to Leslie Webster in 1932, who further disseminated them to various academic and commercial breeders. The Swiss albino mice are descendants of outbred Swiss mice, known for their genetic diversity (Chia et al., 2005). In 1935, Swiss mice were selectively inbred for specific phenotypes over the next 40 years (∼150 generations), leading to the creation of the FVB strain (Rice and O’Brien, 1980). Considerable genetic information has been amassed from research on mouse genetics, oncology, and toxicology, shedding light on the differences between large collections of inbred *Mus musculus* populations now available with various breeders. Future systematic investigations into the genetic divergence and reproductive modes among inbred strains is necessary to obtain valuable insights into the mechanisms driving the disruptive evolution of oocyte’s haploid parthenogenetic blastocyst fitness observed in FVB mice.

In the light of our observations, coupled with the extensive evidence demonstrating the rapid and repeated alterations in reproductive strategies across the animal kingdom, as influenced by environmental factors (Morgan-Richards et al., 2019), it is plausible to speculate that present-day mammalian oocyte reproductive modes and fitness continues to be molded by ongoing selection pressures. This suggests that rare mammals could potentially evolve towards complete asexual reproductive fitness, whether in the wild, captivity, or controlled settings, contingent upon how evolving oocytes navigate evolutionary reproductive constraints, such as those involving extraembryonic placental development, which are currently offset by sperm factors in mammals (Strain et al., 1995).

On early cleavage dynamics, recent scientific endeavors have shed new light on the dynamic and often error-prone nature of embryonic cleavage, particularly evident in the early stages of human embryo development (Currie et al., 2022). The timing and precision of these cleavage events have emerged as critical factors influencing the developmental potential of embryos, underscoring the intricacies inherent in the process of embryogenesis (Fenwick et al., 2002; Lechniak et al., 2008; Wong et al., 2010). Aberrations such as irregular cleavage patterns and compromised cytoplasmic movement have been identified as contributing factors to distorted embryonic division and proliferation during the critical 2-cell-stage (Ezoe et al., 2019; Ohata et al., 2019; Yang et al., 2015).

Blastocysts originating from 1PN zygotes post IVF and ICSI displayed 22.0% mosaic, 3.4% aneuploidy, and no haploidy. Conversely, arrested cleavage-stage embryos demonstrated 28.4% mosaic, 7.4% aneuploidy, and 30.5% haploidy, as determined through FISH (Liao et al., 2009).

In this regard, our investigation of early cleavage dynamics led to the introduction of a novel classification system termed FC-states. This system demonstrated a notable inverse correlation between FC-state and the timeline for exiting totipotency. Additionally, it provided a precise predictive model for evaluating blastocyst generation efficiency and quality in both mouse and human embryos. We did not evaluate the genetic determinants of this phenomenon. Recent studies are shedding light on genetic (transposable elements and transcription factors) regulation of 2-cell blastomeres exiting totipotency (Vega-Sendino et al., 2024). Therefore, future research should investigate mechanisms of enigmatic self-correcting totipotency clock as observed in all our experiments using mouse and human embryos.

In summary, our findings represent a significant advancement in our understanding of embryonic development and embryogenetic blastocyst fitness. By unraveling the intricacies of oocyte development and embryonic cleavage dynamics, we pave the way for future investigations into the evolutionary mechanisms driving reproductive success and genetic diversity maintenance.

The significance of FVB acquiring this blastocyst fitness lies in its ability to challenge conventional evolutionary assumptions. Through this study, we found that haploid blastocysts in the FVB strain exhibit fitness levels comparable to those of diploid blastocysts from sexual reproduction. However, it is noteworthy that there is no evidence suggesting long-term fitness advantages for haploid blastocysts, particularly when compared to diploid parthenogenetic embryos. Despite the normal appearance of diploid parthenogenetic blastocysts, imprinting continues to limit parthenogenetic development in mammals. These findings indicate that the FVB strain may serve as a valuable model for further exploration of the factors influencing reproductive strategies and the evolutionary dynamics between asexual and sexual reproduction. This prompts a reexamination of the mechanisms driving reproductive fitness and adaptation across anisogamous animal populations.

## Acknowledgements

D.N.C. acknowledges support from the United States-India Educational Foundation’s Fulbright-Nehru Doctoral Research (FNDR) Fellowship (mentored by N.K.) and the Government of India’s DST-INSPIRE Fellowship (mentored by G.K.). N.K. received support from the Mayo-NCI SPORE for Ovarian Cancer [CA136393-11CEP] research grant. We thank Dr. Janet Rossant of the Gairdner Foundation and the Hospital for Sick Children, Ontario, Canada, for her valuable review and suggestions for the manuscript.

## Author Contributions

D.N.C., G.K., and N.K. designed the project, and G.K. and N.K. were jointly responsible for its overall execution and quality control. D.N.C. performed all the experiments in N.K. lab. J.F. and T.T. provided technical assistance. G.K. in India, and Y.Z. and N.K. in US provisioned mice for the study. Y.Z. provided clinical data. S.W.L. analyzed clinical data. D.N.C., G.K., and N.K. wrote the manuscript. All authors contributed to the interpretation of the results and read and approved the manuscript.

## Declaration of interests

All authors declare no competing interests.

## Supplementary Figure Legends

**Supplementary Figure 1:**

**A)** The embryo generation efficiency through *in vitro* fertilization, diploid parthenogenetic activation and haploid parthenogenetic activation of oocytes derived from Swiss albino and FVB strains of mice. **B)** Proportion of Dip-AsexE and Hap-ASexE at various developmental-stages, per 100 SexE in Swiss albino strain of mice. **C)** Proportion of Dip-AsexE and Hap-ASexE at various developmental-stages, per 100 SexE in FVB strain of mice. **D)** Schematics illustrating embryo development from syngamy/PN fading up to 8-cell-stage, with annotations of timelines (tPNf, t1, t2, t3, t4, t5, t6, t7, t8), embryonic cell cycle durations (ECC1, ECC2, ECC3), and durations for blastomere synchrony (S2 and S3). Durations (in hours) of **(E)** embryonic cell cycle 1 (ECC1); **(F)** embryonic cell cycle 2 (ECC2); **(G)** embryonic cell cycle 3 (ECC3); **(H)** blastomere synchrony during 2nd cell cycle (S2); and **(I)** blastomere synchrony during 3rd cell cycle (S3)-for SexE, Dip-AsexE and Hap-ASexE derived from Swiss albino and FVB strains. The inter-strain data in panels E-I were analyzed using two-way ANOVA-Sidak’s multiple comparison test, *** p<0.001, ** p<0.01, * p<0.05; **(J)** Representative images of blastocysts from SexE, Dip-AsexE and Hap-ASexE of Swiss albino and FVB mice.

**Supplementary Figure 2:**

**A)** Incidence of FC-states in all embryos derived from Swiss albino and FVB strains of mice. **B)** OR plot depicting effect of FC-states on the probability of SexE, Dip-AsexE and Hap-ASexE derived from Swiss albino and FVB strains of mice to achieve developmental success to the blastocyst-stage (OR below 1) or to undergo arrest in earlier stages of preimplantation embryo development (OR above 1). The values are normalized to the mean success rate of all the mice embryos combined. Effect of FC-states on the duration (in hours) of embryo development from early-to-proper-2C-stage in murine **(C)** SexE, **(D)** Dip-AsexE, and **(E)** Hap-ASexE. Effect of FC-states on the duration (in hours) of embryo development from proper 2C-to-4C-stage in murine **(F)** SexE, **(G)** Dip-AsexE, and **(H)** Hap-ASexE.

**Supplementary Figure 3:**

**A)** Correlation of developmental time between FC-state (early-to-proper-2C) and exit of totipotency (proper-2C-to-4C) in successful (formed blastocyst) or arrested embryos derived from Swiss albino (**i**) SexE (**ii**) Dip-AsexE (**iii**) Hap-ASexE, and FVB (**iv**) SexE, (**v**) Dip-AsexE and (**vi**) Hap-ASexE. Linear regression analysis of timelapse data was performed for data in panel A. p-value<0.05 was considered significant, and the graph was shaded green. P-values>0.05 were considered non-significant (ns) and the graphs were shaded red. Effect of FC-states on the duration (in hours) of embryo development from sexual/asexual fertilization to PNf in murine **(B)** SexE, **(C)** Dip-AsexE, and **(D)** Hap-ASexE. Effect of FC-states on the duration (in hours) of embryo development from PNf to early-2C-stage in murine **(E)** SexE, **(F)** Dip-AsexE, and **(G)** Hap-ASexE.

**Supplementary Figure 4:**

Effect of FC-states on the duration (in hours) of Human-SexE development from **A)** early-to-proper-2C-stage, **(B)** proper 2C-to-4C-stage, **(C)** time of fertilization to time of PNf, and **(D)** time of PNf to early-2C-stage. Effect of FC-states of Human-SexE embryos on the grade of **E)** Inner cell mass and **(F)** trophectoderm, scored by Gardner grade, calculated retrospectively. The data in panels A-F is analyzed using one-way ANOVA-Tukey’s multiple comparison test, ** p<0.01; **G)** Effect of FC-states in Human-SexE, on the decision to use blastocysts for embryo transfer in patients, calculated retrospectively. The groups were compared using Fisher’s exact test, **** p<0.0001, *** p<0.001, * p<0.05. Correlation of developmental time between FC-state (early-to-proper-2C) and exit of totipotency (proper-2C-to-4C) in Human-SexE embryos that formed **(H)** properly-differentiated inner cell mass (ICM), and embryos that got arrested or formed poorly-differentiated ICM, **(I)** good trophectoderm (TE), and embryos that got arrested or formed poor TE, and **(J)** embryos that formed good quality blastocysts with properly differentiated ICM and TE or poor quality blastocysts/arrested embryos. Linear regression analysis of timelapse data was performed for data in panels H, I and J. p-value<0.05 was considered significant, and the graph was shaded green. p-values>0.05 were considered non-significant (ns) and the graphs was shaded red. **K)** OR plot depicting effect of FC-states on the probability of Human-SexE, to achieve developmental success by forming good quality blastocysts (OR below 1) or to undergo arrest in earlier stages of preimplantation embryo development/ form poor quality blastocysts (OR above 1). The values are normalized to the mean success rate of all the human embryos combined.

**Supplementary Video 1:** Time-lapse imaging showing the dynamics of FC-I in murine embryos.

**Supplementary Video 2:** Time-lapse imaging showing the dynamics of FC-II in murine embryos.

**Supplementary Video 3:** Time-lapse imaging showing the dynamics of FC-III in murine embryos.

**Supplementary Video 4:** Time-lapse imaging showing the dynamics of FC-IV in murine embryos.

**Supplementary Video 5:** Time-lapse imaging showing the dynamics of FC-I in human embryos.

**Supplementary Video 6:** Time-lapse imaging showing the dynamics of FC-II in human embryos.

**Supplementary Video 7:** Time-lapse imaging showing the dynamics of FC-III in human embryos.

**Supplementary Video 8:** Time-lapse imaging showing the dynamics of FC-IV in human embryos.

**Supplementary Table 1:** Cross-species and cross-strain data of blastocyst rate (%) in Hap-AsexE and Dip-AsexE in comparison to SexE from our data, and from literature.

**Supplementary Table 2:** Cross-species and cross-strain data of blastocyst rate (%) in Dip-AsexE in comparison to SexE from our data, and from literature.

## Methods

### Mus musculus strains

Adult female and male Swiss albino mice (8-12 weeks) were obtained from Jackson Laboratory (JAX stock #034608) and maintained in standard mouse facility at Mayo Clinic, MN, USA or obtained from the breeding colony at Central Animal Research Facility, Manipal Academy of Higher Education, Manipal, India. The FVB mice (8-12 weeks) from the in-house breeding colony were also maintained at the mouse facility at Mayo clinic, MN, USA. The mice were fed with standard chow diet and water *ad libitum* and maintained under 12h-12h light-dark cycle at 23 ± 2°C. All procedures were reviewed and approved by the Institutional Animal Ethics Committee of Kasturba Medical College, Manipal and the Mayo Clinic Institutional Animal Care and Use Committee (IACUC).

### Murine Oocyte Collection

The adult female mice were stimulated using 5 IU of pregnant mare serum gonadotropin (PMSG) and 10 IU of human chorionic gonadotropin (hCG) at an interval of 48 h. At 13.5 h post hCG administration, the mice were dissected, and the oocyte cumulus complexes (OCCs) were collected by teasing the oviduct in M2 medium. The OCCs were either subjected to *in vitro* fertilization (IVF) to obtain sexually fertilized embryos (SexE) or to parthenogenetic activation to obtain haploid or diploid parthenogenetic embryos (ASexE).

### *In Vitro* Fertilization of Murine Oocytes

The IVF was carried out as described in (Crasta et al., 2023). Briefly, the caudal spermatozoa were released in Earle’s balanced salt solution (EBSS) medium containing 0.1% bovine serum albumin (BSA) and incubated at 37°C, 5% CO_2_ and 20% O_2_ for 2 h to induce capacitation. Next, the motile spermatozoa were collected by swim up technique using EBSS medium containing 2.5% BSA. The sperm count was adjusted to 3-5 millions/mL to prepare insemination droplets (80 µL) covered with light mineral oil (9305, Irvine Scientific). OCCs were collected from super-ovulated female mice and randomly transferred to each insemination droplet and incubated at 37°C, 5% CO_2_ and 20% O_2_. At 13 h post insemination, the fertilized oocytes were washed and observed under the inverted microscope (X400) to confirm fertilization and transferred to the embryoscope chamber slide (n=4 embryos/ well) in M16 medium overlaid with light mineral oil, for time-lapse imaging.

### Parthenogenetic Activation of Murine Oocytes

The OCCs were subjected to the haploid activation medium (10 mM strontium chloride in M16 medium without Ca^2+^ and Mg^2+^) or the diploid activation medium (10 mM strontium chloride and 1 µg/mL cytochalasin D in M16 medium without Ca^2+^ and Mg^2+^) overlaid with light mineral oil for 3 h (Crasta et al., 2022) at 37°C, 5% CO_2_ and 20% O_2_. This was followed by denuding using 0.75 mg/mL of hyaluronidase, wash using M16 medium, and observation under the inverted microscope (X400) to confirm activation, which was indicated by the presence of 1 PN for haploid parthenotes and 2 PN for diploid parthenotes. These were then transferred to the embryoscope chamber slide (n=4 embryos/ well) in M16 medium overlaid with light mineral oil, for time-lapse imaging.

### Clonally Time-lapse Tracked Blastocyst Generation Assay

Four embryos were cultured per well in an EmbryoSlide (Vitrolife) containing 25µL of M16 medium per well overlaid with 1.3 mL light mineral oil. The EmbryoSlide was placed in an EmbryoScope ES-D time-lapse (Vitrolife) incubator (at 37°C, 5% CO_2_ and 5% O_2_) for time-lapse imaging. Precise developmental kinetics were captured of the developing embryos in 3 planes at 5 min intervals for 120-150 hours, until all the embryos reached the expanded blastocyst-stage. The timings of cell divisions were determined manually from the recordings, and it included timeline for formation of 2-cell (t2), 3-cell (t3), 4-cell (t4), 5-cell (t5), 6-cell (t6), 7-cell (t7), 8-cell (t8), morula (tM) and blastocyst (tB), as well as blastocyst expansion, with time of PN fading (tPNf) taken as 0 and the rest calculated accordingly. Durations of embryonic cell cycles ECC1 (t2-tPNf), ECC2 (t4-t2), ECC3 (t8-t4) and time for blastomere synchrony S2 (t4-t3), S3 (t8-t5) were also determined (Ciray et al., 2014). During the first-cleavage, we further subdivided the timeline into early-2C, irregular-2C, and proper-2C, and the timing of each was documented. Additionally, we have developed first-cleavage (FC)-dynamic-state scoring-system (FC-I to -IV) based on irregular cleavage-axis, delayed cytokinesis, irregular shaped blastomeres, and cytoplasmic fragments and each embryo was assigned to FC-states based on scoring criteria.

### Human *In Vitro* Fertilization, Time-Lapse Imaging, and Clinical Data

The study was approved by the Institutional Review Board of Mayo Clinic. Total of 93 autologous embryos from 10 patients who underwent intracytoplasmic sperm injection (ICSI) cycles with oocyte retrieval between January 2014 and January 2023 at Mayo Clinic, MN, USA were randomly selected for analysis. Patients who used donor eggs, or donor sperms and patients who did not consent to the use of their information from the medical records were excluded from the study. After oocyte retrieval, oocytes were inseminated by ICSI for fertilization (referred to as the human sexually fertilized embryos (Human-SexE) in this study), and placed individually in EmbryoScope+ time-lapse system (Vitrolife, Sweden) for culture at 5% O_2_ and 5.8% CO_2_, and imaged at 9 different planes at 15 min intervals. The embryo time-lapse imaging was assessed retrospectively, and the time from insemination to pronuclear fading (PNf), time of division to 2-cell-stage (2C), and time from 2-cell-stage to 4-cell-stage were recorded (Ciray et al., 2014). During the first-cleavage, we further subdivided the timeline into early-2C, irregular-2C, and proper-2C, and the timing of each was documented. Embryos which underwent direct cleavage to 3 or more cells without 2-cell-stage were excluded from analysis. Each embryo that underwent first cleavage to form 2-cell stage was assigned FC-states based on the scoring criteria established in mice. In addition to time-lapse data and FC-states scoring, each embryo was assessed for blastulation, conventional morphologic evaluation of blastocyst quality (modified Gardner Grading system) (Pierson et al., 2023), and the fate of the embryo was decided based on score (trasnferable or not). Scores 1 and 2 were considered as good quality blastocysts with properly-differentiated inner cell mass and trophectoderm. Scores 3 and above were considered as poor quality blastocysts.

### Statistical Analysis

Statistical analysis was performed using the GraphPad Prism 8.0.1 software, California, USA. The two-way Analysis of variance (ANOVA) – Tukey’s multiple comparison test or Sidak’s multiple comparison test were used to compare various groups. The percentage data was analysed using GraphPad Instat 3 software, California, USA. The Fisher’s exact test were used to compare the various groups. Correlation analysis was done using linear regression analysis and the R^2^ values and p-values were noted. Significance level p<0.05 was considered as statistically significant.

## Notes

### Competing Interest Statement

The authors have declared no competing interest.

