## Supplementary figures and images for "Haploid Asexual Blastocyst Fitness Varies Across Mouse Strains Related to Efficiency of Exit From Totipotency"

### Supplementary Figure 1-4

#### Supplementary Figure 1

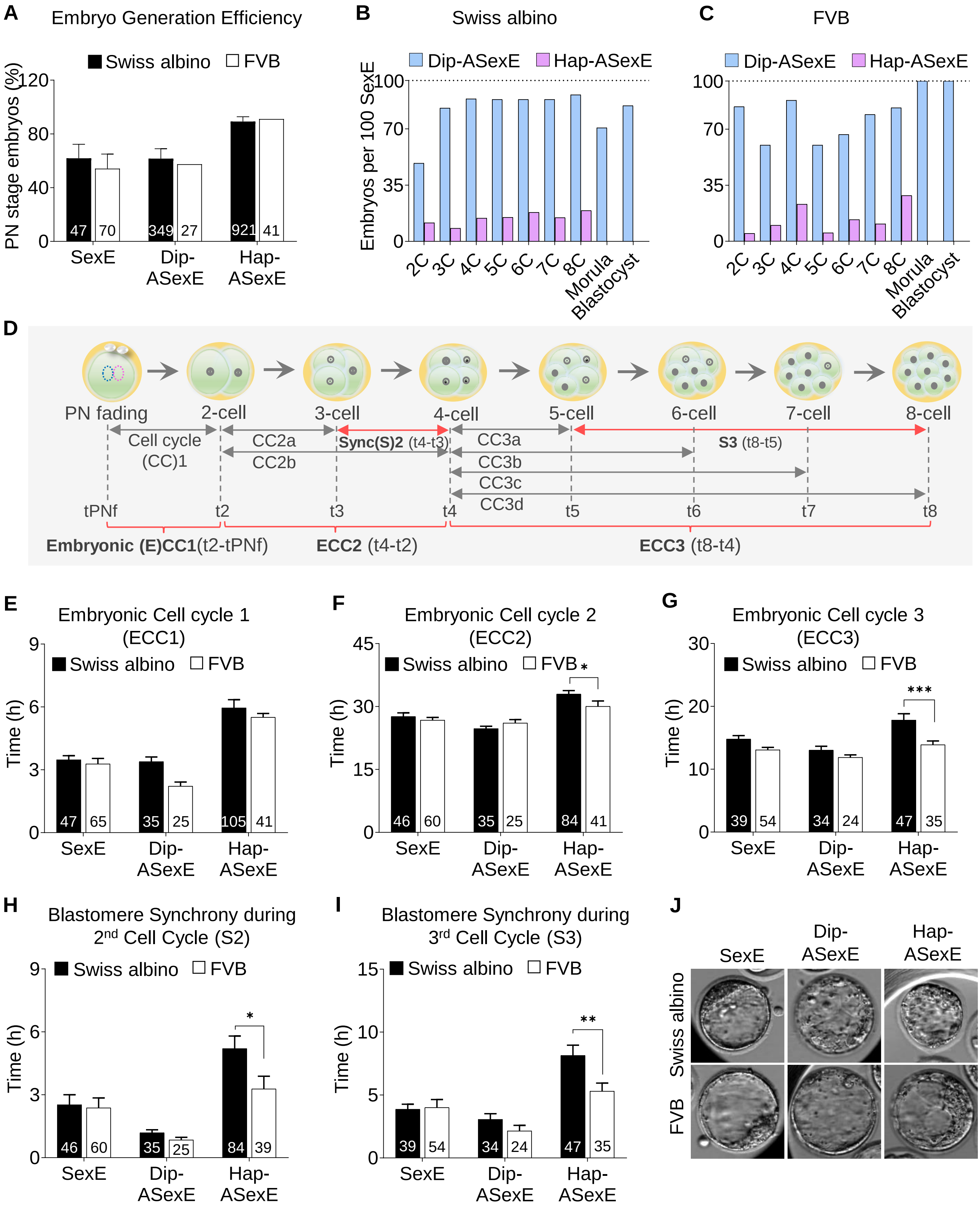

##### Supplementary Figure 2

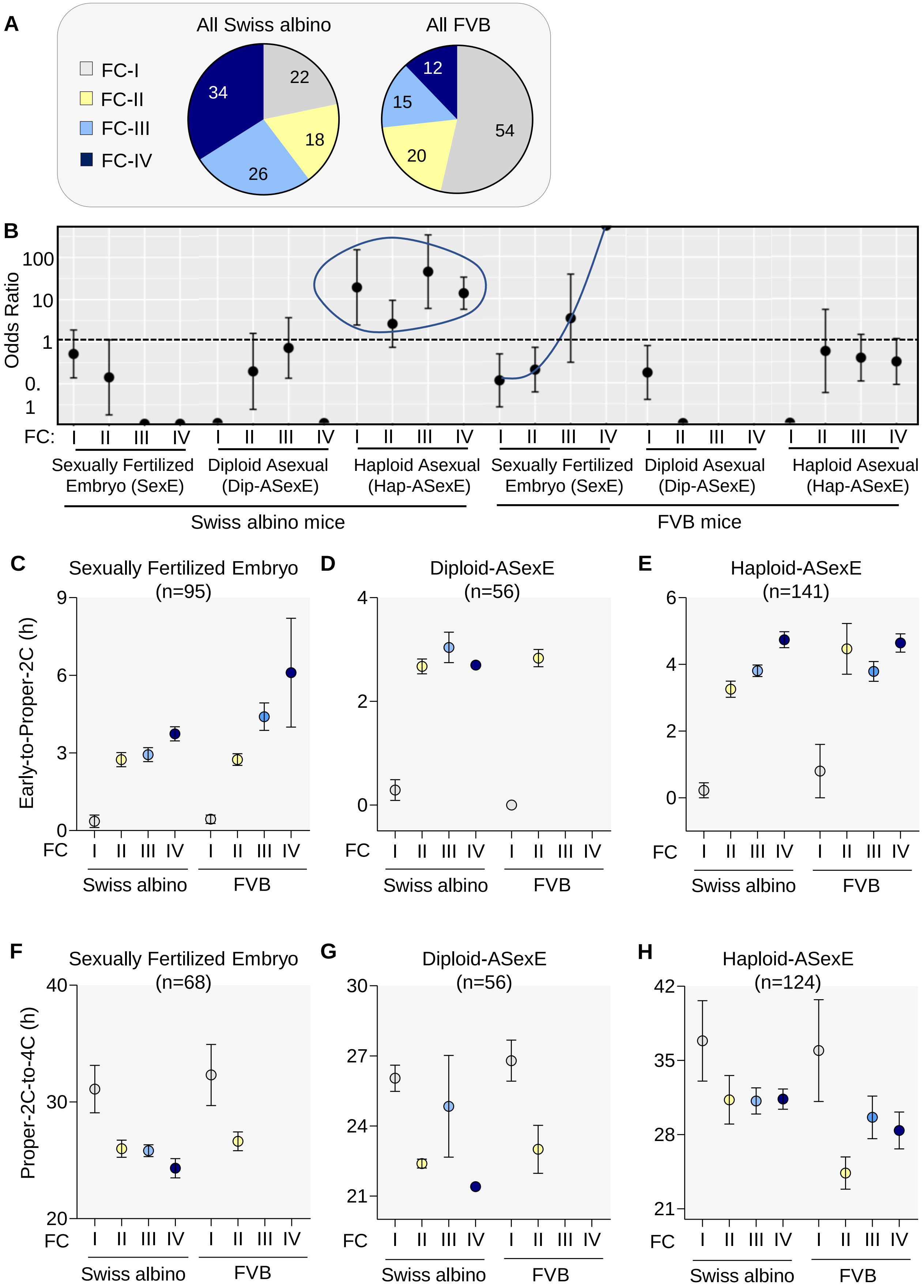

#### Supplementary Figure 3

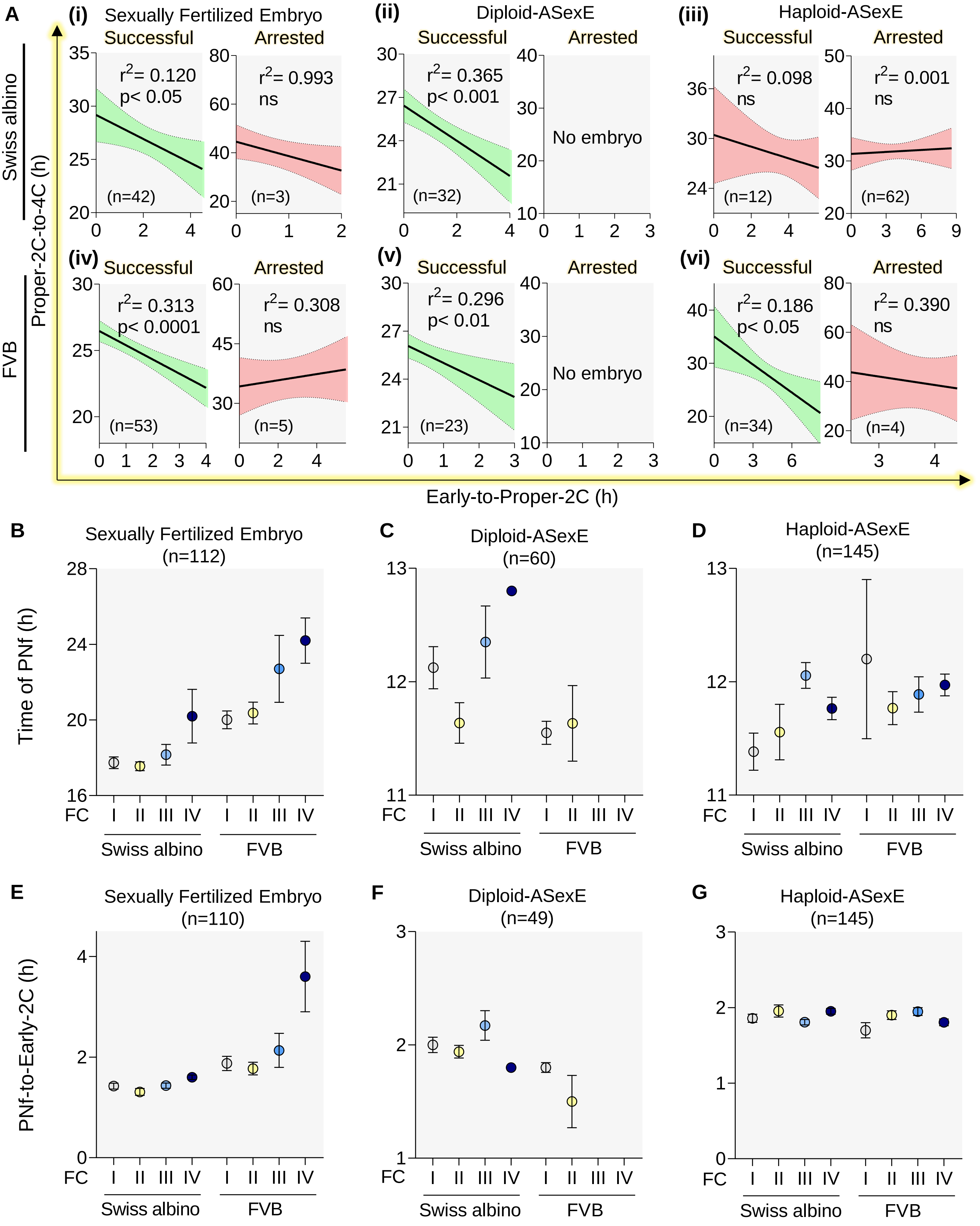

**Supplementary Figure 4**

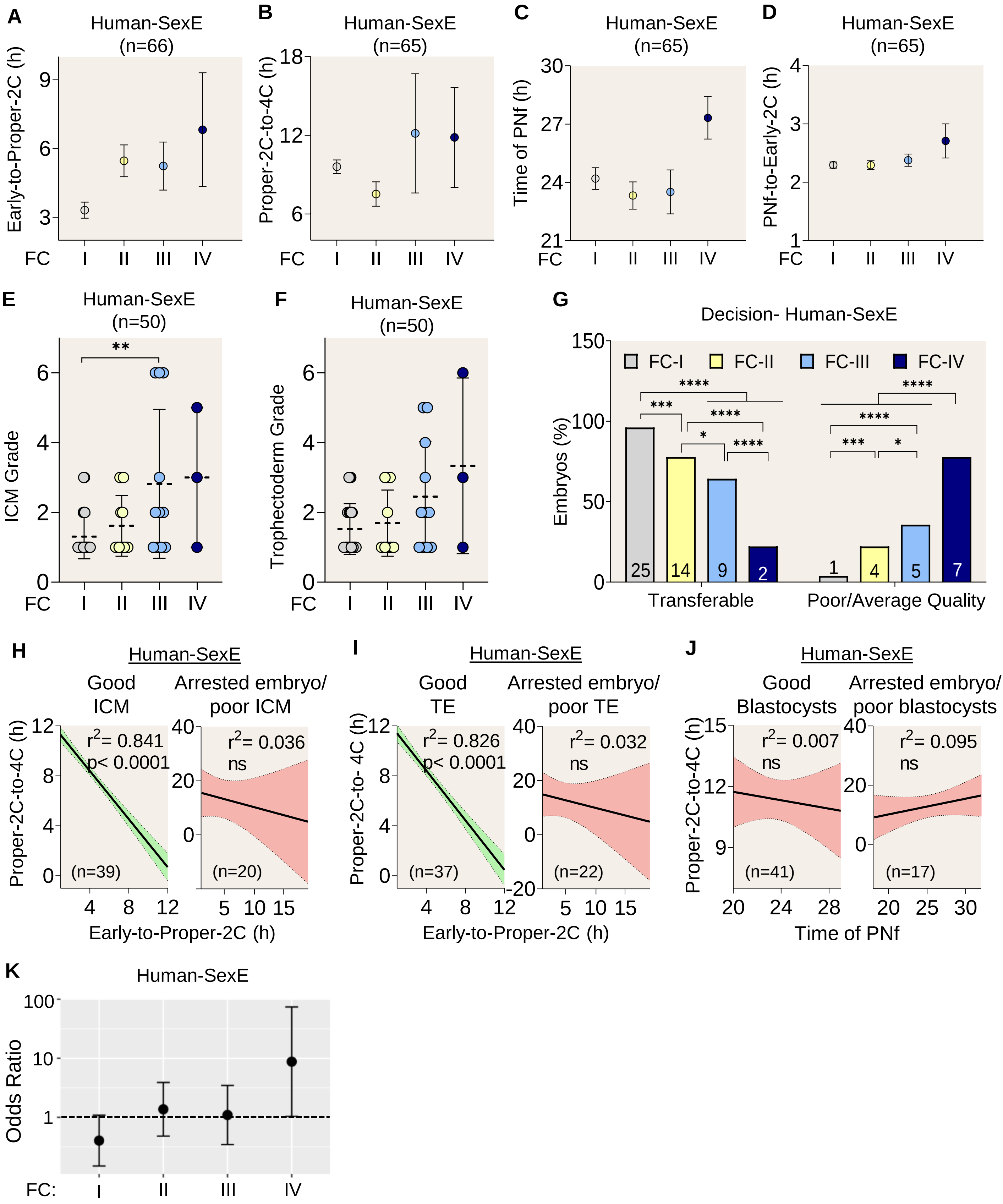
