## Supplementary Table 1-2 for "Haploid Asexual Blastocyst Fitness Varies Across Mouse Strains Related to Efficiency of Exit From Totipotency"

| Species | Sub-strain | %Dip-ASexE over SexE | %Hap-ASexE over SexE | References |
|---|---|---|---|---|
| *Mus musculus* | Swiss albino | 94.24 | 12.10 | Our data |
| *Mus musculus* | FVB | 110.58 | 103.65 | Our data |
| *Mus musculus* | B6C3F1 | 98.86 | 5.68 | Liu et al., 2002 |
| *Bos spp.* | Unknown | 123.23 | 41.08 | Lagutina et. al. 2004 |
| *Bos spp.* | Unknown | 74.43 | 15.06 | Wang et al., 2008 |
| *Rattus norvegicus* | Sprague-Dawley | 55.29 | 9.22 | Roh et al., 2003 |
| *Sus spp.* | Unknown | 122.71 | 40.72 | Park et al., 2011 |

**Supplementary Table 2**

| Species | Sub-strain | %Dip-ASexE over SexE | References |
|---|---|---|---|
| *Mus musculus* | Swiss albino | 94.24 | Our data |
| *Mus musculus* | Swiss albino | 90.00 | Our data |
| *Mus musculus* | FVB | 110.58 | Our data |
| *Mus musculus* | B6C3F1 | 98.86 | Liu et al., 2002 |
| *Bos spp.* | Unknown | 123.23 | Lagutina et. al. 2004 |
| *Bos spp.* | Unknown | 74.43 | Wang et al., 2008 |
| *Bos spp.* | Unknown | 91.43 | Dinnyés et al., 2000 |
| *Bos spp.* | Unknown | 66.67 | Cevik et al., 2009 |
| *Sus spp.* | Unknown | 104.92 | Gupta et al., 2009 |
| *Sus spp.* | Unknown | 122.71 | Park et al., 2011 |
